# Host-secreted lactate during respiratory viral infection diminishes macrophage antibacterial activity through metabolic reprogramming

**DOI:** 10.64898/2026.07.11.737940

**Authors:** Sadia Sultana, Erin Walsh, Jennifer M. Bomberger

## Abstract

In polymicrobial infections, how the host recognizes and responds to pathogens influences which species will persist to cause chronic infections. The human respiratory tract is a common anatomical site for viral-bacterial co-infections, where primary viral infections predispose to secondary bacterial infections, leading to increased morbidity and mortality. Additionally, co-infections are disproportionately prevalent in people with chronic lung diseases, such as chronic obstructive pulmonary disease and cystic fibrosis. We previously reported that primary viral infections and antiviral interferon (IFN) signaling stimulate *Pseudomonas aeruginosa* (PA) biofilm formation on airway epithelial cells (AECs). IFN signaling induces aerobic glycolysis in AECs and generates lactate as a cellular byproduct. Given that innate immune systems play an integral role in co-infection dynamics, we investigated the role of host-secreted metabolites (i.e. lactate) on innate immune cell activity during respiratory co-infections. We found that exposure to the apical secretions from IFNβ-treated AECs significantly compromised macrophage antibacterial activity, with the soluble metabolite lactate playing an important role. Macrophages used monocarboxylate transporters and G-protein receptors to transport and/or sense lactate, respectively, and this exposure to lactate diminished their bacterial-killing activity in a time-exposure dependent manner. Lactate exposure particularly reprogrammed macrophage cellular metabolism towards an anti-inflammatory state by increasing oxidative phosphorylation and fatty acid oxidation. Collectively, these findings provide insight into metabolites as complex regulators of trans-kingdom interactions and epithelial-macrophage crosstalk during respiratory co-infections.

## Introduction

Complex polymicrobial, cross-kingdom interactions are commonly observed in anatomical sites with diverse microbial communities, such as the gastrointestinal and respiratory tracts (1,2). Increasing evidence shows that acute viral infections predispose to the acquisition of secondary bacterial infection in those sites (3–5). While the clinical manifestations of viral infections vary from mild to those that require hospitalization, viral-bacterial co-infections often lead to worse outcomes (2,6,7). Influenza and respiratory syncytial virus (RSV) are common viral culprits associated with bacterial superinfection in the respiratory tract (8,9). Particularly, RSV infection is the major cause of infant bronchiolitis and is associated with the development of chronic *Pseudomonas aeruginosa* (PA) infections and pulmonary exacerbation in people with chronic lung diseases, such as cystic fibrosis (CF) (2,3,10,11). PA is a Gram-negative opportunistic pathogen, and persistent PA lung infections often lead to severe pulmonary exacerbations; yet the detailed mechanism of how acute viral infections augment the acquisition of chronic PA infections has not been fully elucidated.

The host innate immune system is the first line of defense in both cytokine signaling and pathogen clearance, where macrophages play a pivotal role (12,13). Macrophages are highly plastic populations; *(i)* based on the origin, they can be differentiated into tissue-resident or recruited (monocyte-derived) macrophages, and *(ii)* relying on the environmental cues, they can polarize towards anti-inflammatory or pro-inflammatory states (14–16). Generally, a pro-inflammatory state in macrophages is highly phagocytic with antimicrobial and tumoricidal responses, while an anti-inflammatory state promotes macrophages to resolve inflammation, phagocytize apoptotic cells, and promote tissue repair and remodeling. In people with chronic diseases like CF, recruited macrophages are the predominant macrophage populations and their function to clear infection is impaired (17–19), an effect heightened in a co-infection environment due to impaired macrophage clearance of secondary bacterial infections (18,20–22).

Using a viral-bacterial co-culture biofilm model in association with airway epithelial cells (AECs), our lab has previously shown that RSV infection and subsequent antiviral response *(i)* dysregulates iron homeostasis in the airway (2), *(ii)* releases antiviral extracellular vesicles (EVs) to impair epithelial-macrophage crosstalk (21), and *(iii)* drives epithelial metabolic reprogramming (23). During RSV infection, interferon signaling is induced and AEC metabolism shifts toward aerobic glycolysis. In a metabolic shift to aerobic glycolysis, cells increase extracellular glucose uptake, which is quickly converted to ATP and produces lactate as a byproduct (23). This metabolic rearrangement is a hallmark of highly proliferative cells (such as cancer cells) and is commonly known as the Warburg effect (24). Lactate is a complex modulator in the tumor microenvironment and has long been considered a waste product. However, recent studies highlight lactate as an energy source, substrate for post-translational and epigenetic modifications, and signaling molecule of the innate and adaptive immune cells (25,26). A key study in tumor research by Colegio et al. demonstrated that tumor cell-generated lactate can signal to or be transported into the local macrophage population, thereby polarizing them towards an immunosuppressive (anti-inflammatory) state (27).

Our observation of increased aerobic glycolysis and lactate secretion in virus-infected AECs highlights surprising similarities between the metabolic remodeling during viral infection and the metabolic setting of the tumor microenvironment. It also suggests that host-derived lactate in the respiratory milieu can dysregulate immune (i.e., macrophage) responses during co-infection by preemptively skewing them towards a suppressive state, reducing their ability to clear secondary bacterial infections, the mechanistic details of which have not been investigated. Leveraging our polarized respiratory epithelial model (2,21) in this work, we sought to determine whether host-secreted soluble metabolites (i.e. lactate) can alter naïve macrophage physiology in the local co-infected environment. Our study reveals that lactate-driven immunometabolic regulation of macrophages during respiratory viral infection favors the acquisition of secondary bacterial infection.

## Results

### Host-secreted metabolites drive macrophage dysfunction during respiratory viral-bacterial co-infections

A hallmark of viral-bacterial co-infections is antiviral signaling leading to impaired antibacterial defense, where macrophage dysfunction plays a vital role (1,11,23). We investigated if antiviral interferon signaling in the respiratory epithelium triggers the secretion of factors that impair macrophage antibacterial activity. Previously, our lab showed that primary viral infections and the subsequent antiviral interferon response reprograms airway epithelial metabolism, leading to the secretion of host metabolites in the respiratory milieu (23). To investigate the role of host-secreted metabolites during primary viral infection in regulating antibacterial activity of innate immune cells, we treated macrophages with respiratory epithelial secretions and evaluated macrophage bacterial killing efficiency using an antibiotic-killing assay (21,28). Polarized respiratory epithelial cells CFBE41o- (hereafter AECs) were treated with interferon-β (IFNβ) for 18 hrs to mimic the host antiviral response, and apical secretions were collected (Fig. 1A). Monocytes isolated from the peripheral blood mononuclear cells of healthy human donors were cultured with M-CSF for 8 days to generate monocyte-derived macrophages (MDMs). MDMs were incubated for 60 min with AEC apical secretions, then challenged with the respiratory pathogen *Pseudomonas aeruginosa* (MPAO1) at a multiplicity of infection of 10. Using bacterial update and survival CFUs (colony forming units) (*see Methodology*) we next evaluated MDMs bacterial killing efficiency. We observed that MDMs incubated with the apical secretions from IFNβ-treated AECs are significantly impaired in MPAO1 killing (Fig. 1B). Previously, our lab has reported that antiviral interferon signaling triggers AEC metabolic reprogramming and produces metabolic byproducts, such as lactate, which are secreted into the apical milieu (23). In the current study, we asked if these secreted metabolites can alter macrophage antibacterial activity. We purified the soluble metabolites from interferon-treated AECs and incubated with macrophages before performing the antibiotic killing assay. Interestingly, we observed a similar reduction in MDM killing efficiency when incubated with the secreted metabolites (soluble supernatant fraction) from IFNβ-treated AECs (Fig. 1C). Together, these results are consistent with the conclusion that secretory soluble metabolites in the apical milieu during primary respiratory viral infection can regulate macrophage anti-bacterial activity.

**Figure 1:**
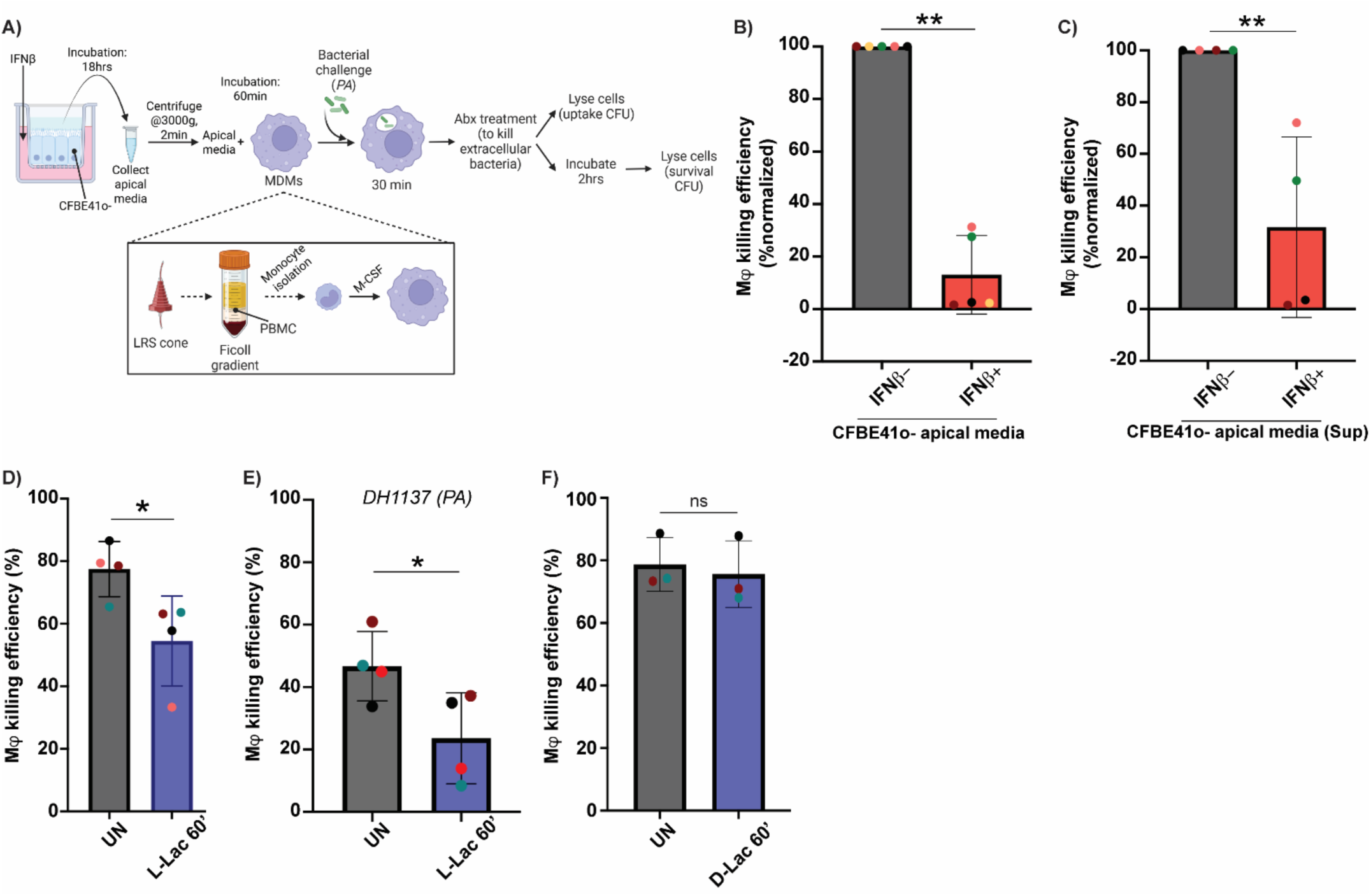
Host-secreted metabolites drive macrophage dysfunction during respiratory co-infections. (A) Experimental workflow to generate monocyte-derived macrophages (MDMs), to collect CFBE41o-apical media after IFNβ treatment, and antibiotic killing assay using respiratory pathogen MPAO1. Assay result displayed as macrophage (Mφ) killing efficiency (B); data are normalized to the control condition (apical media without IFNβ treatment). (C) Apical media from A were ultracentrifuged to separate the soluble supernatant (Sup) fraction and used to analyze Mφ killing efficiency. Antibiotic killing assay after Mφ were pre-treated with (D) 10 mM L-lactate, and (F) 10 mM D-lactate for 60 minutes. Each color denotes a donor. (E) *Pseudomonas aeruginosa* clinical isolate DH1137 was used to analyze macrophage killing efficiency after exposure to 10 mM L-lactate for 60 minutes. Each color denotes a donor. Statistics: linear mixed effects model, donor as a random effect, and treatment as a fixed effect. *p< 0.05, **p< 0.01.

### Host-secreted lactate during viral infection reduces macrophage antibacterial activity

Viral infections are known to trigger metabolic remodeling in AECs, including increased levels of aerobic glycolysis (23,29,30). L-lactate is a by-product of aerobic glycolysis and is present in the infection microenvironment (23,24,26,31–33). It is well established in the cancer biology literature that L-lactate present in the tumor microenvironment can act as a signaling molecule and/or energy source for neighboring immune cells and skew macrophage polarization towards an anti-inflammatory state (24,25,34,35). Therefore, we sought to explore whether lactate present in apical secretions of virus-infected AECs contributes to MDMs’ reduced bacterial-killing efficiency. We incubated MDMs with 10 mM L-lactate, which approximates the lactate concentration during respiratory viral infection (23) and performed the antibiotic-killing assay. MDMs bacterial-killing efficiency was significantly reduced after 60 min incubation with lactate (Fig. 1D), but not after 24 hrs incubation (Fig. S1A); suggesting that MDMs exhibit a time-dependent response to lactate. To determine whether the observed phenotype is a bacterial strain-specific phenotype, we performed the assay using cystic fibrosis lung PA isolate DH1137 (36). Consistent with the findings with PAO1 strain of PA, MDMs showed similar defects in DH1137 killing efficiency (Fig. 1E). Lactate can exist in two isomeric forms, L-lactate and D-lactate; L-lactate is mostly produced by host cells and D-lactate by bacterial species, while some *Lactobacillus* species and *Staphylococcus aureus* can generate L-lactate (37,38). To investigate if MDMs respond similarly to D-lactate, we carried out antibiotic-killing assays after D-lactate exposure, and the data showed no reduction in MDMs killing efficiency (Fig. 1F). Finally, we exposed MDMs to a high concentration of L-lactate that resembles the lactate level in the tumor microenvironment (39) and performed the antibiotic-killing assay to examine if accumulation of lactate can exhibit a similar response. Interestingly, we found that exposure to 30 mM L-lactate for 60 min did not skew macrophage antibacterial activity (Fig. S1B). Together, our experiments support the conclusion that L-lactate (hereafter referred to lactate) negatively impacts macrophage anti-bacterial activity in an exposure time and dose-dependent manner.

### Macrophages exposed to lactate show a defect in bacterial killing

Macrophage efficiency in response to an infection often relies on bacterial engulfment, phagocytic killing, and its activation. Based on the findings that lactate treatment reduced macrophage antibacterial activity, we asked whether this phenotype was due to decrease in bacterial uptake, killing, or both (40,41). Evaluation of bacterial uptake and survival CFU counts suggests a deficiency in MDMs’ bacterial-killing activity rather than in bacterial uptake (Fig. 2A, Fig. S2A). Additionally, we performed macrophage uptake assays using fluorescently tagged *(i)* latex beads and *(ii) Pseudomonas aeruginosa* strain PAO1-tdTomato. Macrophages were treated with lactate, challenged with latex beads or PAO1-tdTomato for 30 min, and analyzed by flow cytometry to quantify the bead- or bacteria-positive macrophage population. Our data showed similar levels of beads or bacterial uptake in both the lactate treated and untreated conditions (Fig. 2B, S2B, S2C). In summary, these findings demonstrate that lactate impairs macrophage bacterial clearance efficacy rather than pathogen uptake.

**Figure 2:**
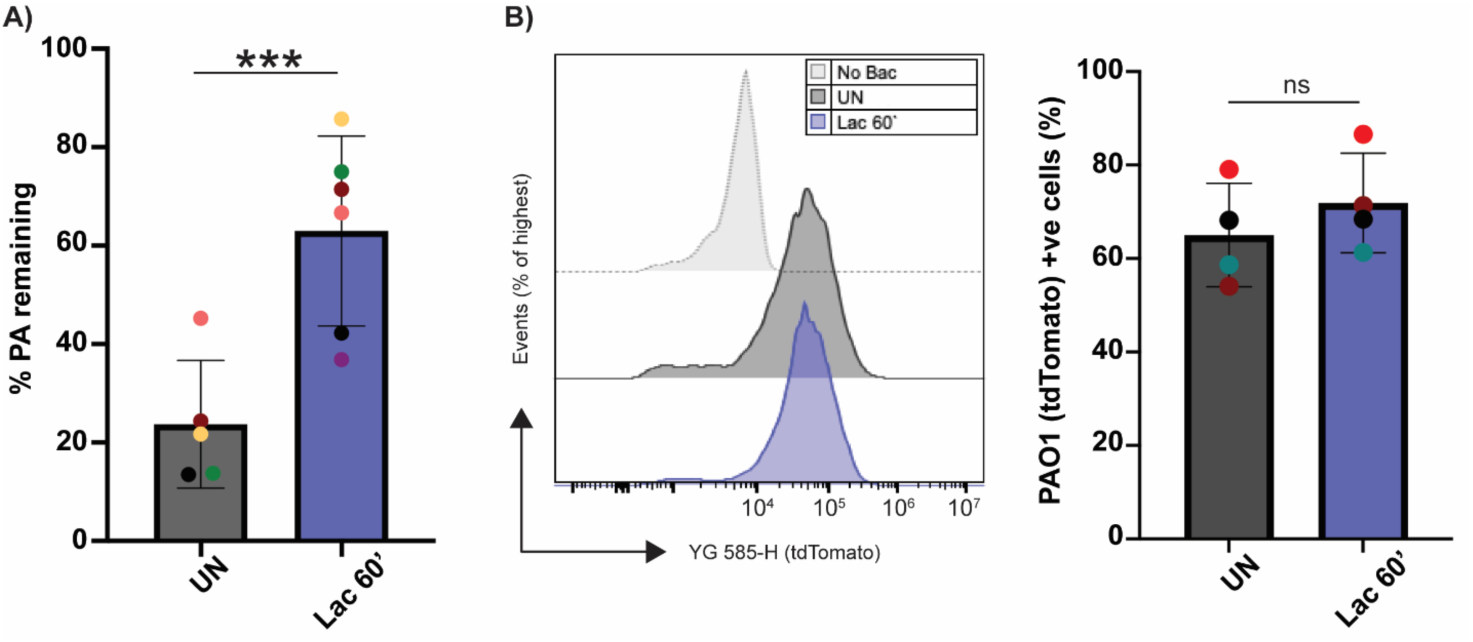
Lactate diminishes macrophage anti-bacterial activity. (A) Mφ MPAO1 uptake and survival counts were used to quantify %PA remaining {(CFU survival/ CFU uptake) × 100%}. (B) Histogram of fluorescent PAO1-tdTomato uptake by Mφ in lactate pre-treated and untreated conditions. Mφ without bacterial exposure was used as a control. The bar graph shows the percentage of Mφ cells positive for PAO1-tdTomato. At least 10000 events were analyzed per condition using the CytoFlex flow cytometer. Each color denotes a donor. Statistics: linear mixed effects model, donor as a random effect, and treatment as a fixed effect. ***p< 0.001.

### Lactate present in the co-infection setting signals through pathways similar to those in the tumor microenvironment

The conventional view of metabolites as merely a “fuel or energy” source to support metabolic pathways has been challenged over the last decade. It is now established that many metabolites can function as intracellular or extracellular signaling molecules to regulate the activity of different enzymes and cellular pathways (42–44). In hypoxic or acidic environments, such as in tumor cells, lactate is transported into the cell via the monocarboxylate transporters MCT1 and MCT4 (35,45) and also sensed by membrane receptors (G-protein-coupled receptors; GPR81/GPR132) (46,47). To test whether lactate present in co-infection settings signals through similar GPCRs and MCTs, we performed the antibiotic-killing assay using specific GPCR and MCT inhibitors (Fig. 3A), as reported in literature (33,48). Pertussis toxin (PTX), a bacterial AB5 toxin, inhibits Gαi/o G protein signaling cascades (49–51), pathway used by the GPR81 and GPR132 for intracellular signaling (46,47). We incubated macrophages with PTX for 18 hrs, followed by 60 min lactate treatment and MPAO1 challenge (Fig. 3A). The presence or absence of PTX did not affect macrophage killing efficiency in the lactate-free condition; however, in the lactate condition, PTX restored macrophage killing efficiency to levels similar to those in the untreated condition (Fig. 3B). Consistent with this idea, we also treated macrophages with 2-cyano-3-(4-hydroxyphenyl)-2-propenoic acid (CHC), an inhibitor of MCT1 and MCT4. Combined treatment with CHC and lactate also restores macrophage killing efficiency to levels comparable to those of untreated macrophages (Fig. 3C). Together, these data suggest that both GPCR-mediated lactate signaling and MCTs-mediated lactate transport suppress macrophage antibacterial activity during respiratory co-infection, consistent with recent literature observation (33).

**Figure 3:**
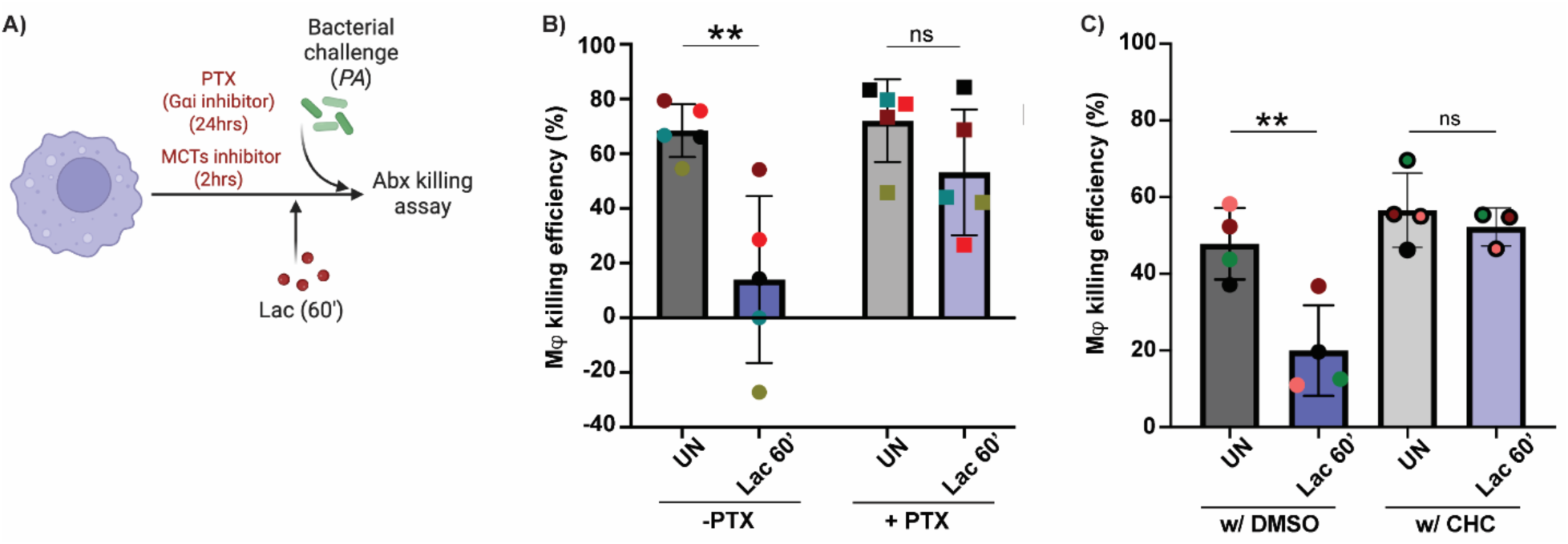
Lactate present in the co-infection setting signals through similar MCTs and G-protein pathways as the tumor microenvironment. Antibiotic killing assay was performed following treatment with G**α**i pathway inhibitor pertussis toxin (PTX, 100 nM) for 24 hr, or MCT-1 and MCT-4 inhibitor *α*-Cyano-4-hydroxycinnamic acid (CHC, 50 nM) for 2 hr (A). Data visualized as Mφ killing efficiency, both (B) PTX and (C) CHC incubation restore Mφ killing efficiency to the untreated level. Each color denotes a donor. Statistics: linear mixed effects model, donor as a random effect, and treatment as a fixed effect. *p< 0.05, **p< 0.01.

### Brief exposure to lactate does not influence macrophage canonical pro-inflammatory pathways

Next, we sought to characterize the interaction dynamics between lactate and macrophages and assessed the physiological features of macrophages after lactate exposure. At first, to trace the possible cytotoxic effect of lactate on macrophages, we performed the glucose-6-phosphate dehydrogenase release assay (52), and found that 10 mM lactate exposure is not cytotoxic to macrophages at either short or long time points (Fig. S3A). Macrophages defective in antibacterial activity are generally referred to as anti-inflammatory macrophages and often exhibit elongated morphology (47). Microscopic analysis of lactate-treated macrophages showed morphological features similar to those of untreated macrophages (Fig. S2C), suggesting that the negative effect of lactate on macrophage bacterial clearance activity is not dependent on morphological changes.

To investigate the classical anti-inflammatory features of macrophages after lactate exposure, we analyzed the changes in cytokine secretion, cellular marker expression, and antimicrobial oxidant production. However, it is important to acknowledge that, while macrophages are generally classified as pro-inflammatory (highly microbicidal) and anti-inflammatory (resolving) states, *in vivo* they are highly plastic populations that exhibit functional/ physiological diversity in response to local environmental cues (14,53–55). MDMs were treated with lactate, supernatants were collected at different time points during MPAO1 incubation, and the cytokine levels were analyzed using DuoSet ELISA. In both untreated and lactate treated conditions, pro-inflammatory cytokine TNFα and anti-inflammatory cytokine IL-10 level remained similar following 30 min and 2 hrs MPAO1 infection (Fig. 4A). Next, we evaluated the expression of anti-inflammatory intracellular (Arg1) and cell surface (CD163, CD206) markers after 60 min of lactate treatment. Macrophages were stained with marker-specific antibodies and flow-cytometrically quantified for the positive cell population. Consistent with the cytokine data, no changes in cell markers were found between lactate-treated and untreated macrophages (Fig. 4B). Reactive oxygen species (ROS) production and phagolysosomal acidification are important features of macrophage bacterial-clearance efficacy (56,57); therefore, we quantified the ROS level and lysosomal pH of lactate pre-exposed MDMs. We stained MPAO1-infected macrophages with CM-H2DCFDA (to detect ROS) and LysoTracker (to detect lysosomes), and flow cytometrically analyzed for positive cell population. The mean fluorescence intensity of the reporter dyes was similar for both the lactate-treated and untreated conditions (Fig. 4C, 4D). Taken together, these data suggest that lactate impairs macrophages’ antibacterial activity, though it does not induce changes in conventional polarization state-specific features-a further confirmation of dynamic plasticity exists in macrophages.

**Figure 4:**
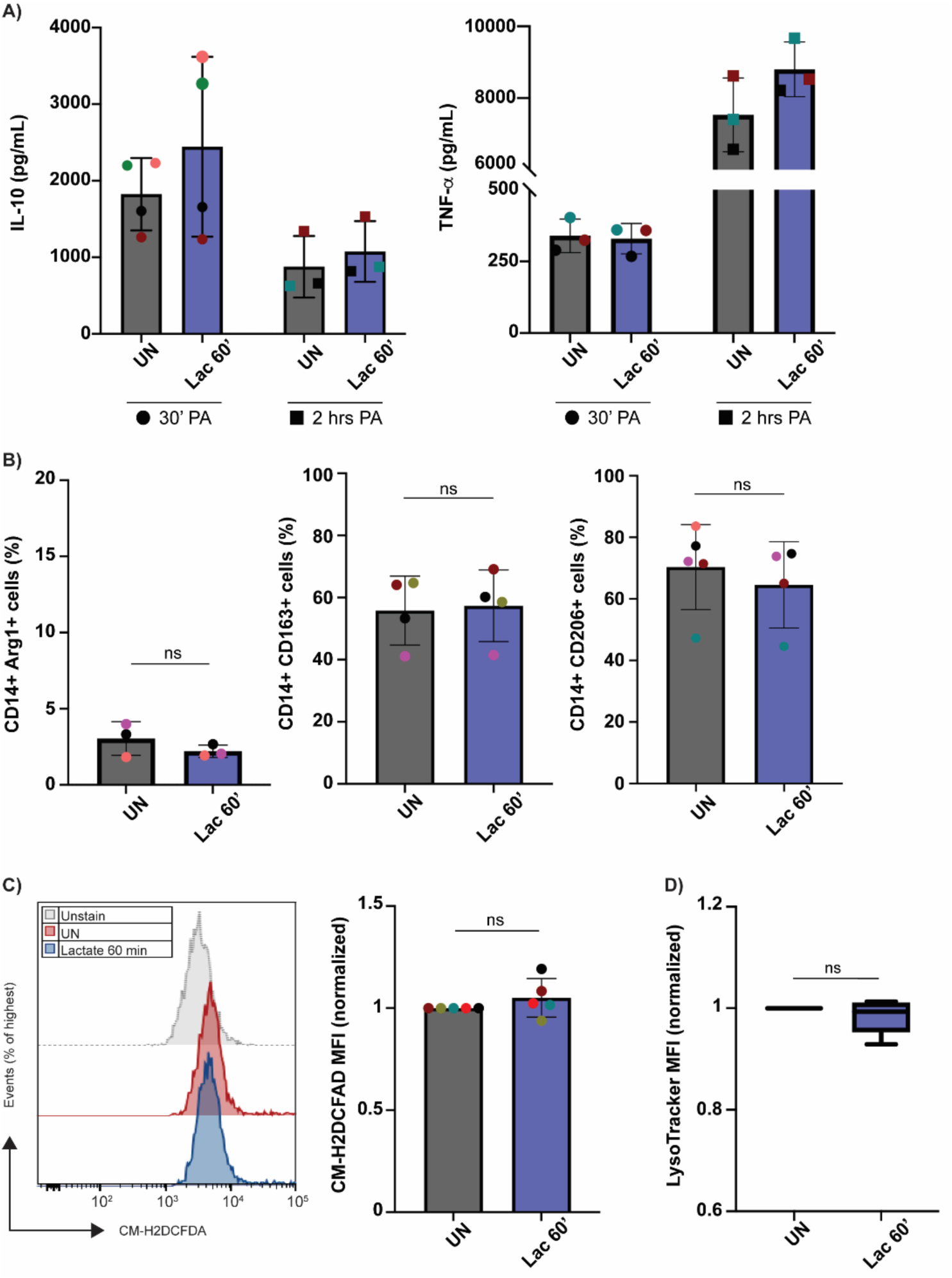
Lactate does not impact macrophage canonical pro-inflammatory pathways. (A) Mφ with or without 10 mM lactate pre-treatment were challenged with MPAO1 for 30 minutes or 2 hr, and supernatant was collected for cytokine analysis using DuoSet ELISA kit. IL-10 and TNFα levels were quantified using the manufacturer-supplied standards. (B) Mφ cell surface (CD163 and CD206) and intracellular (Arg1) markers expression in control or 10 mM lactate pre-treatment condition were analyzed after incubation with specific fluorescent antibody for 30 minutes. Extracellular CD14 was used as an Mφ-specific marker. At least 10000 events were analyzed per condition using the CytoFlex flow cytometer, and Mφ positive for CD14 and respective markers were analyzed using FlowJo software. Mφ were pre-incubated with 10 mM lactate, challenged with MPAO1, stained with (C) CM-H2DCFDA for ROS, or (D) lysotracker-DeepRed for lysosomal pH, and analyzed using CytoFlex flow cytometer. At least 10000 events were analyzed per condition. (C) The bar graph shows the mean fluorescent intensity (MFI) of lactate-pre-treated Mφ, normalized to the untreated control. The histogram shows a representative data set. (D) MFI was calculated after normalization to the untreated control. Each color denotes a donor. Statistics: linear mixed effects model, donor as a random effect, and treatment as a fixed effect.

### Lactate modulates macrophage cellular metabolism

Emerging studies are now highlighting the interconnection between macrophage metabolism, polarization states, and subsequent functional activity in the infection environment (55,58–60). Pro-inflammatory macrophages shift their metabolism towards aerobic glycolysis upon activation, and their TCA cycle is also attenuated to facilitate increased ROS production. In contrast, anti-inflammatory macrophages mostly rely on oxidative phosphorylation (OXPHOS) or fatty acid oxidation (FAO) for their energy source (61). Therefore, we next investigated macrophage metabolic state in the presence of lactate. To test as markers for metabolic changes, we performed mitochondrial oxygen consumption and fatty acid oxidation assays.

To evaluate the real-time metabolic state of macrophages after lactate exposure, we performed mitochondrial stress test using the fluorescence-based MitoXpress Xtra oxygen consumption rate (OCR) and Seahorse assays. For MitoXpress Xtra OCR assay, MDMs were treated with lactate for 30, 60 min, or 24 hrs before obtaining the time-resolved fluorescence reading for 90 min. MDMs OCR was significantly increased after shorter (30/ 60 min) lactate exposure, but not after longer (24 hrs) incubation (Fig. 5A, Fig. S4). This finding is consistent with our previous antibiotic-killing assay results, in which we observed a similar time-course of macrophage antibacterial activity (Fig. 1D, Fig. S1A). To further investigate the macrophage metabolic profile, we conducted a Seahorse assay using the XFe96 Bioanalyzer after sequential addition of specific modulators. Assay data revealed that the difference in OCR after lactate exposure is primarily driven by an increase in basal oxygen consumption and proton leak, while the maximum respiration remained unaffected (Fig. 5B).

**Figure 5:**
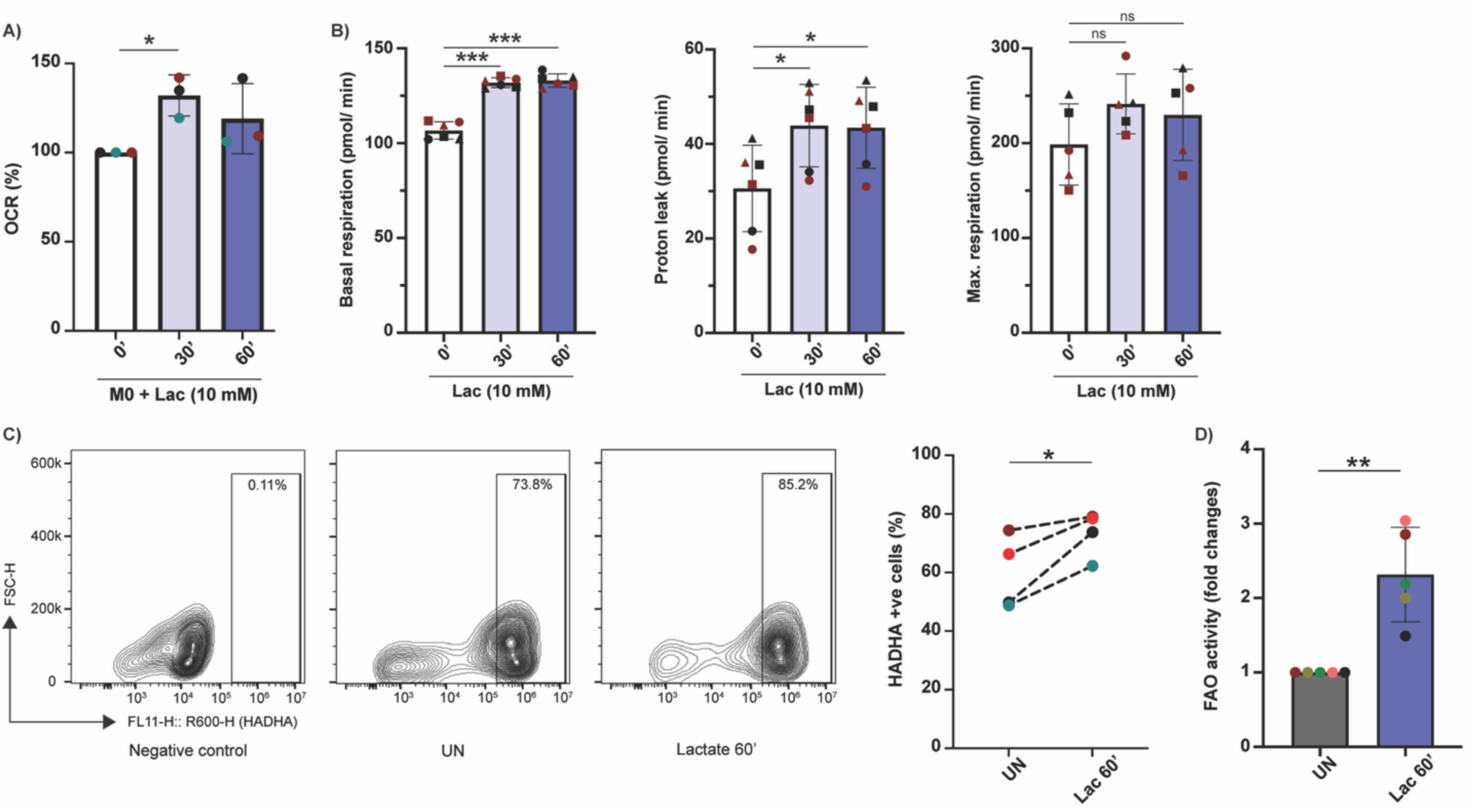
Lactate modulates macrophage cellular metabolism towards an anti-inflammatory state. (A) Mφ oxygen consumption rate (OCR) was spectrophotometrically quantified using MitoXpress Xtra probe after exposure to 10 mM of lactate for 30 and 60 minutes. (B) Mφ was treated with 10 mM lactate and basal respiration, proton leak and maximum respiration was quantified using Seahorse XF Cell Mito Stress test. Each color denotes a donor and shape represents technical replicates for each donor. (C) Mφ was treated with lactate, and the mitochondrial fatty acid β-oxidation pathway enzyme HADHA level was quantified using flow cytometry. Contour plot of representative datasets, and the graph shows Mφ population positive for HADHA. At least 10000 events were analyzed per condition. (D) Fatty acid oxidation (FAO) activity of Mφ cell lysates, with or without lactate pre-treatment, was quantitatively measured using octanoyl-CoA substrate (*see methodology for details*). Each color denotes a donor. Statistics: linear mixed effects model, donor as a random effect, and treatment as a fixed effect. *p< 0.05, **p< 0.01, ***p< 0.001.

Anti-inflammatory macrophages also exhibit changes in lipid metabolism with an increase in mitochondrial fatty acid β-oxidation (FAO) (62,63). Long-chain fatty acids are imported into the mitochondrial membrane, where they are oxidized by mitochondrial dehydrogenases, such as ACADVL and HADHA/B, generating acetyl-CoA, NADH, or FADH2; these fuels the TCA cycle and the electron transport chain to generate ATP (55,64). Therefore, as a secondary confirmation of macrophage metabolic remodeling after lactate exposure, we analyzed *(i)* the level of expression of a mitochondrial fatty acid β-oxidation pathway enzyme HADHA, and *(ii)* the FAO activity of MDM lysates. Exposure to lactate significantly increases both the HADHA-positive cells and FAO activity compared to the untreated control (Fig. 5C, 5D). Collectively, these findings demonstrate that host-secreted lactate during viral infection facilitates macrophage metabolic remodeling, thereby suppressing their antibacterial efficacy and thus, promoting secondary bacterial infections in the airway environment.

## Discussion

The mechanism by which preceding or concurrent viral infections predispose secondary bacterial infections is complex and multifaceted. Viral infections can subvert airway physiology and mucosal immunity, which results in susceptibility to bacterial attachment and failure to control bacterial replication, respectively (8). These defects in the airway environment are exaggerated in people with chronic respiratory conditions, such as cystic fibrosis (CF) or chronic obstructive pulmonary disorder (COPD) (17,18,20). Macrophages, the first line of host innate immune defense, play a decisive role in the co-infection environment by orchestrating cytokine and chemokine responses. Using a specialized *in vitro* model of well-differentiated airway epithelial cells (AECs), we previously reported that antiviral response shifts AECs metabolism towards aerobic glycolysis, resulting in increased secretion of byproducts (i.e., L-lactate) in the apical milieu (23). This metabolic rearrangement also facilitates secondary bacterial infection in the airway by promoting biofilm formation (23); however, the role of the AECs metabolic rearrangement in the airway immune response was not known. In this work, we demonstrated that AECs-secreted soluble metabolites during antiviral response impair macrophage antibacterial activity (Fig 1). We further showed that one of the secretory metabolites, L-lactate, is pivotal to this defective physiology of macrophages by increasing macrophage oxidative phosphorylation and mitochondrial fatty acid oxidation (Fig 6).

**Figure 6:**
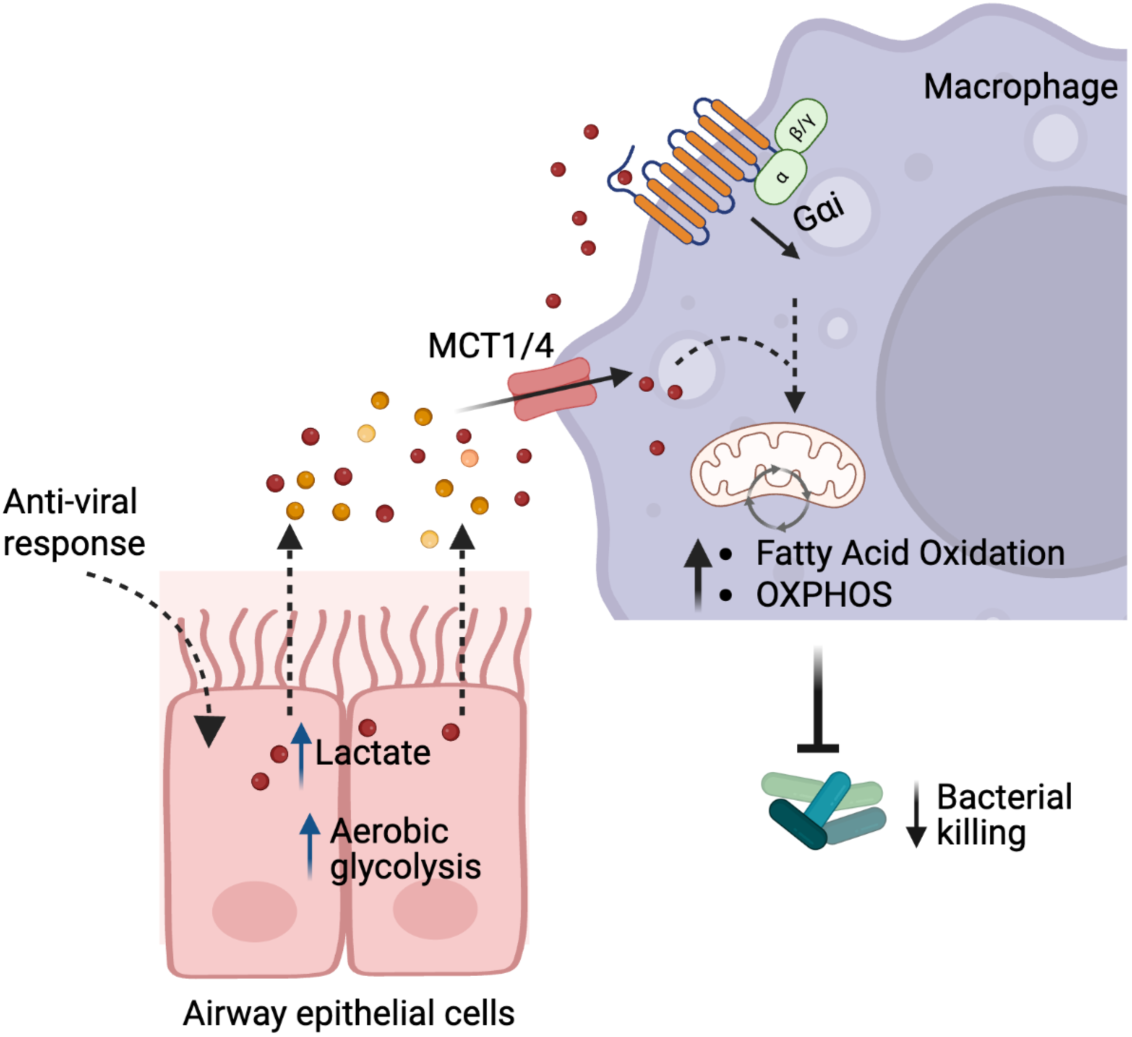
Proposed model: Airway antiviral response triggers elevated levels of lactate generation and subsequent secretion in the apical milieu, which is then transported (by monocarboxylase transporters, MCTs) and/ or sensed (by G-protein receptors) by macrophages recruited in the environment. This lactate transport/sensing facilitates macrophage metabolic reprogramming towards an anti-inflammatory state; ultimately, reducing their antibacterial efficacy.

Viral infections and antiviral responses enhance host cells’ glucose metabolism through aerobic glycolysis, resembling the widely studied Warburg Effect in cancer (45,65). Aerobic glycolysis supports rapid generation of ATP to accommodate the cellular energy demand and it also funnels metabolites to other pathways, including the pentose phosphate pathway (PPP) for nucleosides, fatty acids, and antioxidant synthesis (23,66,67). A hallmark feature of the Warburg effect is the increased production and secretion of lactate into the extracellular environment, which has historically been considered as a waste product. However, growing evidence confirms that lactate modulates immune responses in cancer (27,68–70) and other infection environments (33,71); suggesting that lactate could be one of the critical molecules that link metabolism and immunity. One of such examples is the muscle tissue after exercise, where tissue resident macrophages consume lactate produced by endothelial cells and polarize into an anti-inflammatory state (72). A parallel phenomenon has been observed in the tumor microenvironment, where tumor cell secrete lactate as a means to “hijack” the surrounding macrophage population, leading to an immune shielding of the tumor (73). In addition, some studies have linked respiratory viral infections and higher levels of L-lactate with pulmonary exacerbation, including worsened bacterial infection (23,31,74), although the precise mechanisms are not known. Collectively, these findings have led us to more broadly consider that macrophage consumption and/or sensing of lactate may facilitate homeostatic and pro-regenerative phenotypes in co-infection environments, which inadvertently contribute to secondary bacterial infections in the airway.

Lactate typically exists as two isomers, L- and D-lactate. L-lactate is the main form found in vertebrates, and D-lactate is mainly synthesized by bacteria under anaerobic conditions (75). In a healthy physiological state, lactate (L-lactate) concentration in blood or tissues is approximately 1.5–3 mM, which can reach over 20 mM during sepsis, 40 mM in the inflamed tissues or tumor microenvironment, and ∼9 mM in CF sputum (76,77). We previously reported that, in the co-infection environment lactate concentration increases up to 10 mM (23). Studies using a broad range of lactate concentrations demonstrated that lactate promotes macrophage anti-inflammation polarization and suppresses pro-inflammatory cytokines production (i.e., TNFα and IL-6) (27,78–80). In line with the existing literature, we also observed that 10 mM L-lactate, but not D-lactate, significantly reduces macrophage killing efficiency (Fig 1), indicating that lactate present in the AECs secretion contribute to macrophage functional deficiencies. Moreover, this reduction in macrophage killing efficiency is a time-dependent response, with shorter incubation (60 min) with lactate having an adverse effect compared to longer incubation (24 hrs) (Fig 1, S1). This is particularly remarkable, given that incubation with high concentration of lactate (30 mM) for shorter times (60 min) did not reduce macrophage killing efficiency (Fig S2). Together, these data support that lactate production and accumulation in the host environment potentially act as a complex timer to balance macrophage pro- and anti-inflammatory responses to maintain tissue homeostasis (71,81).

Studies have established mechanisms of lactate transport and sensing via transporters and/or receptors that affect immune cells’ pro- and anti-inflammatory phenotypes by altering cytokine production or cellular metabolic reprogramming via kinase activation and protein modification (i.e., lactylation) (34). Monocarboxylate transporter (MCT) is a member of the solute carrier 16 transporter family, of which there is a total of 14 subtypes (34,82). Among them, MCT1-4 are proton-dependent transporters involved in bidirectional transport of monocarboxylic acid (82–85). Under physiological conditions, MCT1-4 can shuttle lactate between glycolytic and oxidizing cells; however, due to it’s high affinity to lactate, MCT1 transports lactate according to the transmembrane gradient to maintain an intracellular and extracellular homeostasis (44,86). Conversely, MCT4 is a hypoxia-inducible MCT with low affinity for lactate and is adapted to release lactate from tumor cells (87,88). These MCT-mediated lactate transport contribute to the activation or repression of transcription factor nuclear factor kappa-B (NF-κB), thereby regulating the inflammatory response (33,44,89). Consequently, inhibition of MCTs suppress tumor growth and enhance immunotherapy, leading to the emergence of lactate metabolism-targeting drug development (35,48). We, therefore, incubated macrophages with MCTs inhibitor (non-selective) 2-cyano-3-(4hydroxyphenyl)-2-propenoic acid (CHC) and examined their antibacterial activity with or without lactate exposure. Our data showed that inhibition of MCTs restores macrophage bacterial killing efficiency (Fig 2), suggesting lactate uptake and potential intracellular accumulation are associated with the immunosuppressive effect. In addition to the transporters, two G-protein-coupled cell surface receptors (GPCRs), GPR81 and GPR132, for lactate have been reported (46,47). GPR81 is relatively high in adipose tissue but is expressed in other cell types (80). A broad range of lactate concentrations (0.1-30 mM) can activate GPR81, suggesting the metabolite can function as autocrine and paracrine molecule (90). GPR132 is a member of the pH-sensing GPCR family, exclusively expressed in macrophages and other hematopoietic lineages but is absent in adipocytes or cancer cells (91). GPR81 and GPR132 share amino acid homology and signal through the Gαi pathway (92). Accordingly, we incubated macrophages with Gαi pathway inhibitor pertussis toxin (PTX) and examined their antibacterial activity upon exposure to lactate. We found that inhibition of Gαi pathway also restores macrophage bacterial killing efficiency in the presence of lactate (Fig 2). Together, this observation suggests that both GPR-mediated lactate sensing and MCT-associated lactate transport are important to skew macrophage antibacterial activity; which is similar to the recent report by da Silva et al. that lactate uses both MCT-1 and GPR81 mediated pathways to suppress macrophage NF-κB activity (33).

The anti-inhibitory effect of lactate on LPS-stimulated macrophages has been shown previously by others (79,93). However, all those studies have used longer incubation of macrophages and lactate. When investigating macrophage anti-inflammatory phenotypes, such as cytokines, cell markers, reactive oxygen level or lysosomal pH, we did not observe any significant change of these properties after 60 min of lactate pre-incubation (Fig 4). This observation is not atypical, given that 60 min exposure may not be enough to visualize changes in those physiological properties. A study by Shi et al also supported this, as their RNAseq analysis of BMDMs (bone marrow derived macrophages) stimulated with LPS and lactate for 3 hr showed no significant induction of anti-inflammatory marker genes. Further kinetic experiment by qPCR revealed that inhibition of pro-inflammatory genes by lactate occurs as early as 1 hr, while the induction of anti-inflammatory genes (i.e., Arg1 and Retnla) is a later event (>3–6 hr) (48). Besides these changes, pro- and anti-inflammatory macrophages are also characterized by specific metabolic adaptations to support their activities and polarization in specific contexts. Pro-inflammatory macrophages show increased glycolysis and accumulation of itaconate and succinate due to TCA cycle brakes, leading to Hypoxia Inducible Factor 1α (HIF1α) stabilization and sustained glycolytic metabolism. Conversely, anti-inflammatory macrophages rely on oxidative phosphorylation (OXPHOS), and fatty acid oxidation (FAO), that provide substrates for the TCA cycle and electron transport chain (55). When investigated for these immunometabolic changes, we found that lactate exposure significantly elevates macrophage OXPHOS metabolism, particularly basal respiration, and also induces FAO and expression of enzymes associated with the FAO pathway (Fig 5). Collectively, these findings demonstrate that lactate present in the co-infection environment shifts macrophage polarization towards an anti-inflammatory phenotype by driving metabolic reprogramming (Fig 6).

Our work highlights a new avenue of macrophage-epithelial crosstalk during viral infection; however, we acknowledge that other metabolites in apical secretions may also contribute to macrophage function in the clinical setting. The use of a reliable co-infection model will potentially strengthen our interpretation, which is the focus of future studies in the lab. In sum, our findings highlight a mechanism behind the widespread clinical observation that viral infection predispose the acquisition of secondary bacterial infections, particularly in people with chronic diseases such as CF. By better understanding the mechanistic details of the complex interactions among respiratory pathogens, host metabolites, and immune system, we can refine existing therapeutic targets including the use of metabolic inhibitor therapeutics.

## Methodology

### Cell lines and bacterial strains

Immortalized human CF airway epithelial cell line CFBE41o- (referred to herein as AECs) were cultured in minimal essential media (MEM) supplemented with 10% fetal bovine serum (FBS), 2 mM L-glutamine, 5 U/mL penicillin– 5 μg/mL streptomycin (P/S), and 0.5 mg/mL Plasmocin prophylactic at 37°C in 5% CO_2_ (21,23). Cells were seeded on 12 mm transwell filters (Coster) for 8–10 days at the air-liquid interface. After well polarization/ differentiation, cells were basolaterally treated with 1000 U/mL IFNβ (R&D) for 18 hr at 37°C with 5% CO_2_. For bacterial strains, *Pseudomonas aeruginosa* MPAO1, clinical isolate DH1137 (36), and fluorescent strain PAO1-tdTomato were used. Unless otherwise mentioned, bacteria were grown aerobically at 37°C in LB broth (Lennox, Millipore Sigma).

### Monocyte-derived macrophage isolation

Human monocyte-derived macrophage (MDMs) were generated from de-identified healthy donor peripheral blood mononuclear cells (PBMCs) purchased from the DartLab (Dartmouth Hitchcock Medical Center). PBMCs were isolated from blood LRS cones via a Ficoll-density gradient, and monocytes were negatively selected from PBMCs using the Classical Monocyte Isolation Kit (Miltenyi) (21). Isolated monocytes were seeded in non-tissue culture plates and cultured with 100 ng/mL recombinant human macrophage colony-stimulating factor (rhM-CSF; R&D) in cRPMI media (RPMI-1640 supplemented with 10% FBS, 2 mM GlutaMAX-I, and 100 U/mL penicillin–100 μg/mL streptomycin) for 6-7 days to differentiate into macrophages. Twenty-four hours prior to all treatments (days 6-7 of macrophage culture), the culture media were changed to antibiotic-free cRPMI (RPMI-1640 supplemented with 100 ng/mL rhM-CSF, 10% FBS, and 2 mM GlutaMAX-I).

### Antibiotic killing assay

Macrophage ability to clear bacterial pathogens after lactate pre-treatment was examined using the well-established antibiotic killing assay (21,28). Macrophages were cultured in duplicate per condition in 48-well non-tissue culture plates as described above. 60 min/ 24 hr before the assay, macrophages were treated with 10 mM lactate or left untreated. Next, cells were challenged with bacteria (MOI = 10) and centrifuged at 500× g for 5 min to promote macrophage-bacteria interaction, followed by 30 min incubation at 37°C with 5% CO2. Afterward, cells were incubated for 1 hr with antibiotic cocktail media (RPMI supplemented with 500 μg/mL gentamicin, and 500 U/mL penicillin–500 μg/mL streptomycin) to kill extracellular bacteria. One duplicate well cells were then washed twice with DPBS (−/−), immediately lysed with 0.1% Triton X-100, serially diluted in PBS, and spotted on LB agar plates to measure bacterial uptake CFU. The remaining duplicate well was given low-antibiotic cRPMI media (RPMI supplemented with 10% FBS, 25 μg/mL gentamicin, and 500 U/mL penicillin–500 μg/mL streptomycin) and incubated for an additional 2 hr before being lysed to collect bacterial survival CFUs. Results are calculated as Macrophage killing efficiency (%) [{(Avg. CFU Uptake-Avg. CFU Survival) / Avg. CFU Uptake} ∗ 100]; or % bacteria remaining [(Avg. CFU Survival / Avg. CFU Uptake) ∗ 100].

### Antibiotic killing assay with apical media from AECs

Influence of host-secreted metabolites on macrophage antibacterial activity was examined using an antibiotic killing assay with apical media. AECs were grown as mentioned earlier and basolaterally treated with 1000 U/mL IFNβ (R&D) for 18 hr at 37°C with 5% CO_2_. After 18 hr incubation, the apical media were collected and centrifuged at 3000x g for 3 min to remove cell debris. The collected supernatants were used to treat macrophages for 60 min prior to performing the antibiotic killing assay. To separate extracellular vesicles, the supernatants were again ultracentrifuged at 100,000x g for 60 min using Optima TLX Ultracentrifuge. The soluble fraction was used in the antibiotic-killing assay, and pellet (EV) was solubilized in 900 μL of RPMI and used in the antibiotic-killing assay. Macrophage killing efficiency was calculated as above and normalized to the IFNβ-control.

### Fluorescent PAO1 and latex-bead uptake assay

Macrophage uptake efficacy was analyzed using fluorescently labelled PAO1 or latex-beads. Macrophages were pre-incubated with 10 mM lactate for 60 min, followed by challenge with td-Tomato PAO1 (MOI= 10), or 1 μm latex beads (Sigma) (MOI= 5). Cells were incubated for 30 min at 37°C with 5% CO_2_, washed with PBS (2x), and lifted from the plate using 200 μL cell dissociation buffer (Gibco). After centrifugation for 3 min at 1500 rpm, the cell pellet was resuspended in 200 μL pre-warmed FACS buffer (2% FBS and 1mM EDTA in DPBS −/−) supplemented with 0.25 μg/mL propidium iodide. Cells were incubated in the dark for an additional 15 min at 37°C and 5% CO2, then analyzed using the Beckman Coulter CytoFlex flow cytometer. At least 10000 events were recorded, and data were analyzed using FlowJo 10.10.0 software. For microscopic visualization, macrophages were seeded on uncoated 48-well MatTek plates and visualized under the Nikon Eclipse Ti2 microscope. Representative images are shown and scale bar is 100 μm.

### Antibiotic killing assay using antagonists

Macrophages were incubated with monocarboxylate transporters (MCT1 and MCT4) inhibitor 2-Cyano-3-(4-isopropylphenyl)-2-propenoic acid (CHC), and G-protein signaling Gαi pathway inhibitor pertussis toxin (PTX) to examine the effect of these pathways on lactate transport or sensing, respectively (33,94,95). Macrophages were pre-incubated with 0.5 mM CHC (Sigma) for 1 hr or with 100 ng/mL PTX (Calbiochem) for 17 hr before performing the antibiotic killing assay, as described above.

### Cytokine analysis

Cytokine production by macrophages was analyzed using the Luminex human 48-plex assay. Macrophages were pre-treated with 10 mM lactate for 60 min, followed by 2 hr of incubation with MPAO1. After bacterial incubation, macrophage supernatant was collected for cytokine analysis.

### Cell surface marker expression assessment

Macrophage cell surface markers expression following lactate exposure was analyzed using a flow cytometry-based assay. Macrophages were treated with 10 mM of lactate for the mentioned times, washed with PBS, and lifted from the plate using cell dissociation buffer (Gibco). For extracellular markers (CD-14, CD-163, CD-206), cells were incubated with BD staining solution supplemented with CD14-APC, CD163-PE-cyan, and CD206-eFluro450 (Invitrogen) at a concentration of 2.5 μg/mL, 5 μg/mL, and 5 μg/mL, respectively. For the intracellular marker (Arg1), cells were fixed and permeabilized with Cytofix/CytoPerm (BD Biosciences) according to the manufacturer’s instructions and stained with Arg1-eFluro450 (Invitrogen) antibody at 2 μg/mL. Finally, cells were resuspended in PBS and counted using a Beckman Coulter CytoFlex flow cytometer. FlowJo software was used for data analysis.

### Oxygen consumption assay

Macrophage oxidative phosphorylation (OXPHOS) metabolism was evaluated using the fluorescent MitoXpress oxygen consumption assay (Agilent). 1x e5 cells/ well were seeded in a non-tissue culture 96-well plate and incubated as per the mentioned condition. 30 or 60 minutes prior to the experiment, the media were changed to clear RPMI supplemented with 10 mM lactate. 10 μL MitoXpress reagent was added to each well according to the manufacturer’s instructions, and time-resolved fluorescence was measured in a Synergy Neo2 plate reader at 30 sec intervals for 90 min at 37°C.

### Assessment of ROS production and lysosomal pH

Macrophage ROS generation and lysosomal pH gradient were evaluated using the fluorescent probes 5-(and-6)chloromethyl-2,7-dichlorohydrofluorescein diacetate acetyl ester (CM-H2DCFDA) and Lysotracker-DeepRed, respectively (Invitrogen). Untreated or 10 mM lactate-treated macrophages were challenged with MPAO1 (MOI= 10) for 30 mins, lifted from plates using cell dissociation buffer (Gibco), and incubated in FACS buffer supplemented with 5 μM CM-H2DCFDA for 15 min, or 50 nM Lysotracker-DeepRed for 30 min. Fluorescent signals were assessed using the Beckman Coulter CytoFlex flow cytometer, and data were analyzed with FlowJo software.

### Fatty acid oxidation assay

A flow cytometry-based and a spectrometry-based assay were used to analyze macrophage fatty acid oxidation efficiency after lactate exposure. Flow-cytometry assay (Abcam) was used to examine the abundance of mitochondrial fatty acid β-oxidation pathway enzyme HADHA. Lactate-pre-treated cells were lifted using cell dissociation buffer and fixed with 4% paraformaldehyde. Next, cells were permeabilized and stained with anti-HADHA primary and goat *α*-mouse IgG secondary antibody according to the kit instructions. Finally, cells were analyzed using a Beckman CytoFlex flow cytometer, and data were interpreted using FlowJo software. The colorimetric assay (AssayGiene) was used to quantitatively measure fatty acid oxidation (FAO) in macrophages. Cells were isolated after lactate treatment, lysed, and the soluble protein fraction was collected. The BCA (Sigma) assay was used to quantify protein level, and an equal amount of protein across samples was used to quantify FAO activity as per the manufacturer’s protocol.

### Statistical analysis

Each experiment was performed using at least three randomly selected de-identified donors’ PBMCs. Data are presented as mean ± SD. A linear mixed-effects model was performed in RStudio, with donor as a random effect and treatment as a fixed effect.

---

**Figure S1:**
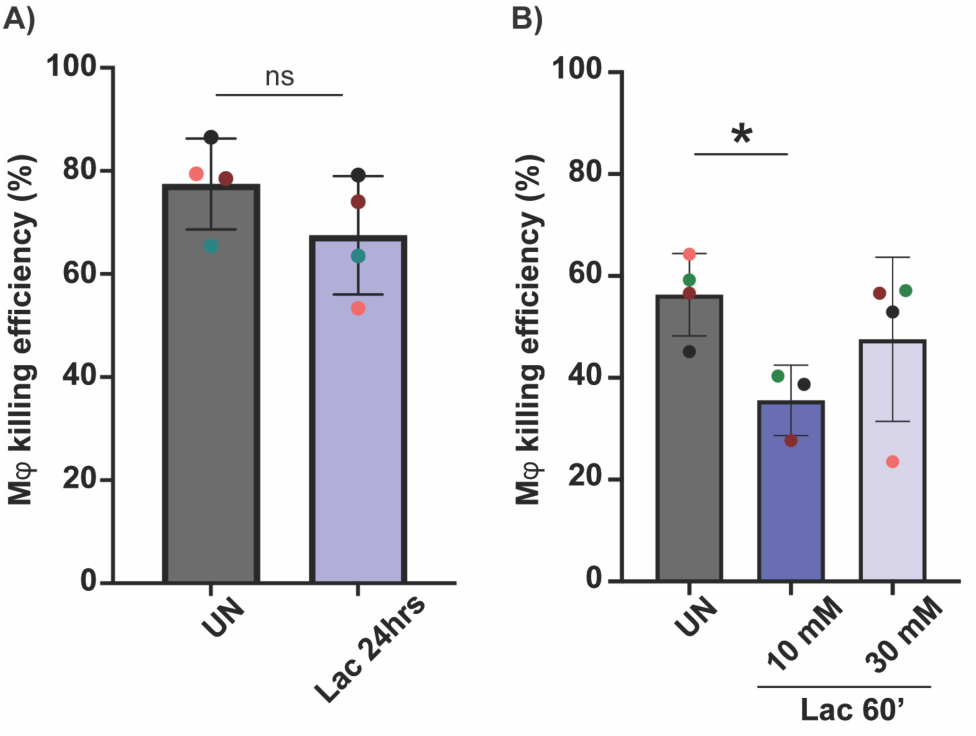
(A) Mφ were exposed to 10 mM lactate for 24 hr, and killing efficiency was analyzed using an antibiotic killing assay. (B) Mφ were treated with 10 mM or 30 mM lactate for 60 min, followed by antibiotic killing assay to analyze macrophage killing efficiency. Each color denotes a donor. Statistics: linear mixed effects model, donor as a random effect, and treatment as a fixed effect. **p< 0.01.

**Figure S2:**
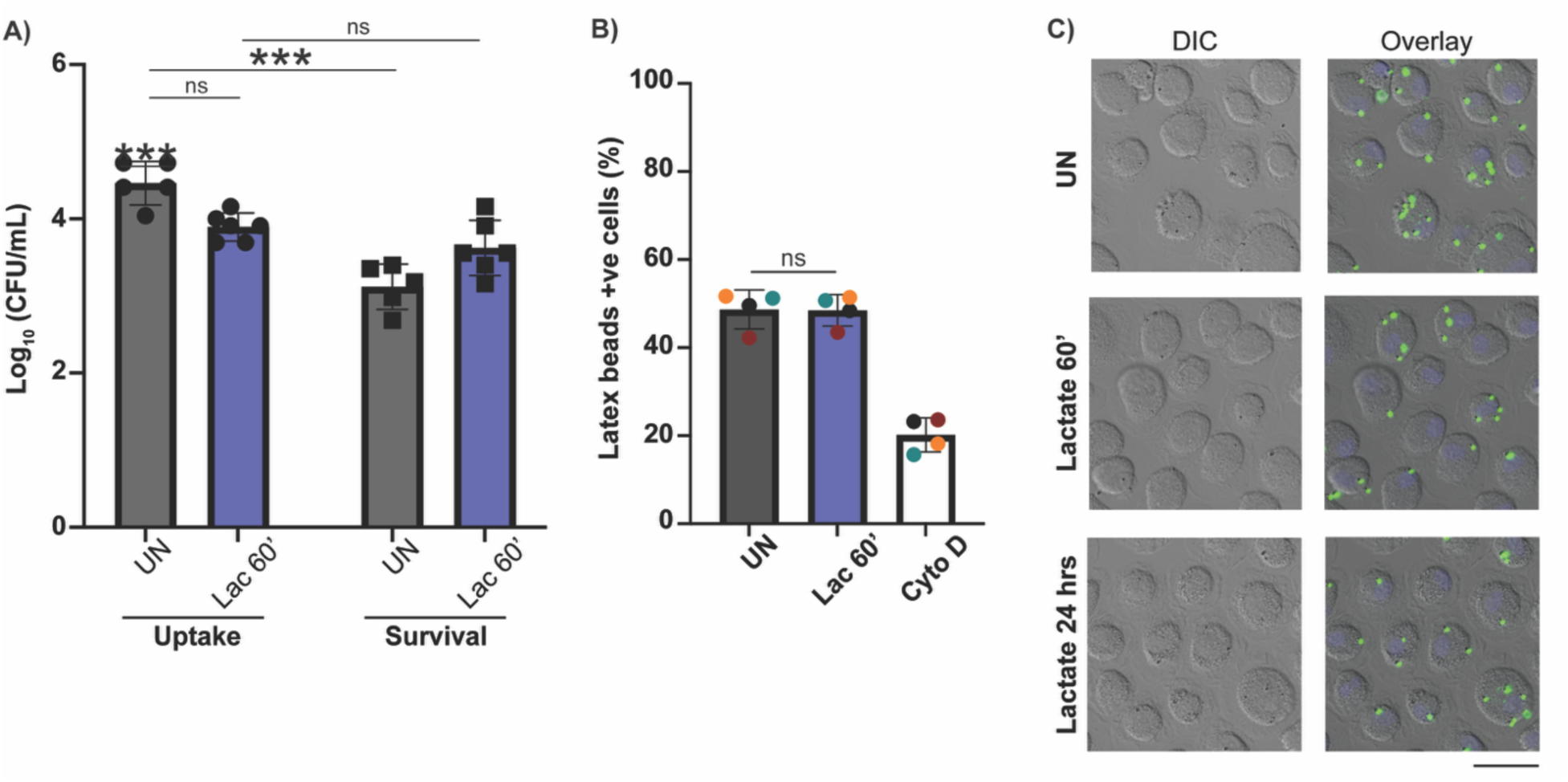
A) Mφ MPAO1 uptake and survival count from the antibiotic killing assay in the lactate treated condition or untreated control. (B) Mφ with or without lactate treatment were incubated with fluorescent 1 μm latex beads at an MOI of 5 for 30 minutes, and bead uptake was analyzed using a Cytoflex flow cytometer. Mφ treated with 10 μM cytochalasin D was used as a negative uptake control. Each color denotes a donor. Statistics: linear mixed effects model, donor as a random effect, and treatment as a fixed effect. (C) Mφ were treated with lactate, exposed to latex beads, fixed using 4% paraformaldehyde, and microscopically visualized. Representative data is shown. Scale bar 100 μm.

**Figure S3:**
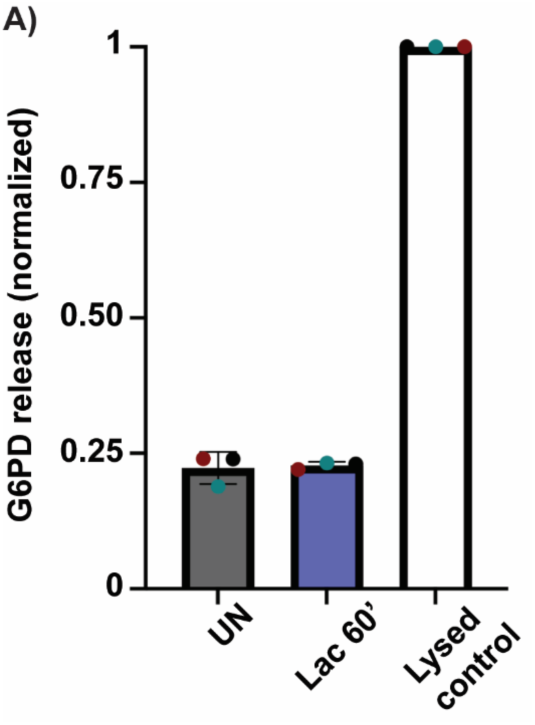
(A) Mφ supernatant was collected following lactate treatment, and glucose-6-phosphate dehydrogenase (G6PD) release was quantified as a measure of cellular cytotoxicity. Each color denotes a donor. Statistics: linear mixed effect model, donor as a random effect, and treatment as a fixed effect.

**Figure S4:**
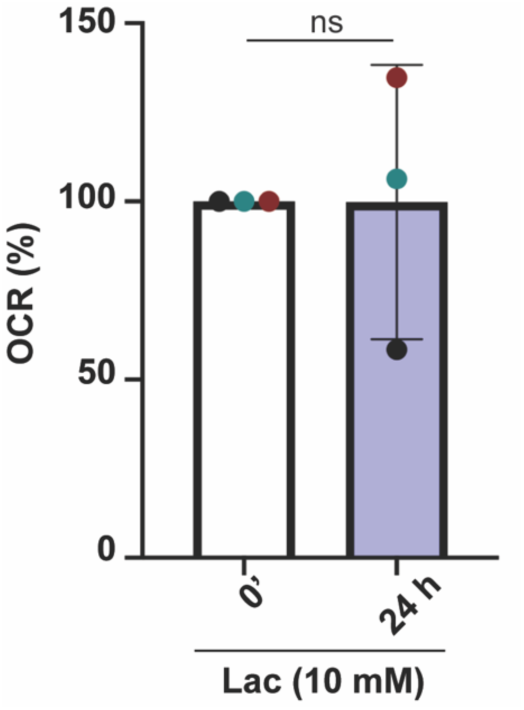
Mφ was exposed to 10 mM of lactate for 24 hrs and oxygen consumption rate (OCR) was spectrophotometrically quantified using MitoXpress Xtra probe. Each color denotes a donor. Statistics: linear mixed effect model, donor as a random effect, and treatment as a fixed effect.

## Acknowledgements.

Research reported in this publication was supported by NIH R01HL169973 (to J.M.B.), Cystic Fibrosis Foundation BOMBER24R0, BOMBER18G0 (to J.M.B.), and SULTAN25F0 (to S.S.). CytoFlex flow cytometry is supported by Dartmouth Cancer Center grant P30CA023108. We thank DartLab at the Dartmouth Hitchcock Medical Center for the LRS cones collection, Dr. Rebecca Valls (Bomberger lab) for statistical analysis support, and Lia Michaels (Bomberger lab) for PAO1-tdTomato strain. Finally, we are grateful to the present and former members of the Bomberger lab for their valuable feedback on this study. The content is solely the responsibility of the authors and does not necessarily represent the funders’ views. Illustrations were generated using BioRender.

## Authors Contributions

Sadia Sultana, Conceptualization, Investigation, Methodology, Data curation, Formal analysis, Visualization, Writing – original draft, Writing – review and editing | Erin Walsh, Investigation, Methodology | Jennifer M. Bomberger, Conceptualization, Funding acquisition, Supervision, Writing – review and editing.

